# Atypical mediofrontal theta oscillations underlying cognitive control in kindergarteners with autism spectrum disorder

**DOI:** 10.1101/2020.12.08.416370

**Authors:** George A. Buzzell, Hannah R. Thomas, Yeo Bi Choi, So Hyun Kim

## Abstract

**Background:** Children with autism spectrum disorder (ASD) often exhibit deficits in cognitive control. Neuroimaging approaches have implicated disruptions to medio-frontal cortex (MFC) structure and function. However, prior work has not directly tested whether young children with ASD exhibit disruptions to task-related theta oscillations thought to arise from the MFC.

**Methods:** Forty-three children with ASD and 24 age- and gender-matched typically developing (TD) peers performed a child-friendly Go/No-go task while 64-channel electroencephalography (EEG) was recorded at kindergarten-entry. Time-frequency approaches were employed to assess the magnitude of mediofrontal theta oscillations immediately following error (vs. correct) responses (“early theta”), as well as later emerging theta oscillations (“late theta”). We tested whether error-related mediofrontal theta oscillations differed as a function of diagnosis (ASD/typical) and timing (early/late theta). Additionally, links to social and academic outcomes were tested.

**Results:** Overall, children showed increased theta power following error vs. correct responses. Compared to TD children, children with ASD exhibited a selective reduction in error-related mediofrontal theta power during the late theta time window. There were no significant group differences for early theta power. Moreover, reduced error-related theta power during the late, but *not* early, time window significantly predicted poorer academic and social skills.

**Conclusions:** Kindergarteners with ASD demonstrated a selective reduction in error-related mediofrontal theta power during a relatively late time window, which is consistent with impairments in specific cognitive processes that recruit top-down control. Targeting these particular cognitive control processes via intervention prior to school-entry may promote more successful functional outcomes for children with ASD.

## Introduction

The ability to self-monitor and flexibly adapt one’s behavior in response to changes in the internal or external environment refers to neurocognitive processes known as *cognitive control* (i.e., executive functions) (1,2). In addition to social communication deficits, a core symptom domain of autism spectrum disorder (ASD; (3), a substantial body of work associates ASD with deficits in cognitive control (4-9). Work in typically developing (TD) children (10), adolescents (11), and adults (12) links cognitive control to a particular pattern of task-related brain oscillations within the theta band (∼4-7 Hz). However, the study of such brain oscillations and their link to cognitive control in children remains limited. Moreover, direct comparisons of task-related theta oscillations between TD and ASD children are largely unexplored. This reflects a crucial gap in the understanding of cognitive control among children with ASD, given that experimental studies have demonstrated such oscillatory activity is causally linked to cognitive control (13,14). Clinically, deficits in cognitive control early in life can have cascading effects on later social and academic outcomes (7,15-19). Thus, an improved understanding of cognitive control neural dynamics in children with ASD is critical for the development of targeted and effective interventions to maximize the outcomes of these individuals. The current study leverages time-frequency analyses of EEG recorded during a cognitive control task to test whether the magnitude of task-related theta oscillations differs in kindergarteners with ASD, compared to TD controls.

Individuals with ASD display deficits in behavioral tasks requiring cognitive control (7,20), including inhibitory control tasks (21). At the neural level, the mediofrontal cortex (MFC)—to include the anterior cingulate cortex (ACC)—is typically activated when healthy individuals perform cognitive control tasks (22). However, decreased blood-oxygen level dependent (BOLD) activity within the MFC (as measured via fMRI) has been observed while individuals with ASD perform inhibitory control tasks (23,24). Moreover, in TD individuals, the MFC is known to become more activated in response to events conveying a need to increase control, such as following error commission (25). However, individuals with ASD also display reduced BOLD activation of the MFC in response to errors, compared to correct responses (26). These findings are consistent with studies of brain structure differences between ASD and TD, which demonstrate differences in the morphometry of MFC subregions, including the ACC (27).

While studies employing fMRI provide evidence for disrupted MFC function in ASD, there is a crucial line of evidence that has gone largely unexplored. In particular, it remains unclear as to whether task-related mediofrontal theta oscillations are disrupted in young children with ASD. Mediofrontal theta is increased during tasks requiring cognitive control, as well as in response to events signaling a need for control, including error commission (10,11,28).

Similarly, mediofrontal theta dynamics observed during cognitive control tasks are thought to be generated, at least partially, within the MFC (29). Crucially, mediofrontal theta oscillations—which can be assessed non-invasively via EEG—do not simply provide an additional metric for assessing cognitive control. Instead, mediofrontal theta reflects a direct readout of a brain mechanism that has been causally implicated in cognitive control (13,14). Theta is thought to serve as an organizing rhythm that supports—through synchronization—a dynamic network of brain regions underlying cognitive control (12,30). Moreover, the enhanced temporal resolution of EEG provides the opportunity to measure the neural dynamics of cognitive control with greater specificity in time. Thus, examining mediofrontal theta in those with/without ASD would provide crucial information regarding the nature of *how* and *when* cognitive control dynamics are disrupted for this clinical population. However, to our knowledge, no direct comparisons of response-related theta in ASD vs. TD in young children have been explored.

While task-related mediofrontal theta has not been directly compared between ASD and TD, comparisons of resting state EEG provides promising evidence for abnormalities within the theta range for ASD (31,32). However, resting state theta power reflects a qualitatively different neural process, functionally and anatomically distinct from task-related mediofrontal theta (33-36). Additionally, in a recent study we found that variability in task-related mediofrontal theta within children with ASD was predictive of academic outcomes (37). However, a key limitation of this work is that we did not include a TD group, preventing any direct comparisons between ASD and TD.

The current study directly tests whether task-related mediofrontal theta oscillations are disrupted in young children with ASD compared to TD. Based on prior work demonstrating that individuals with ASD exhibit impairments on inhibitory control tasks (21), and reduced fMRI-BOLD activation of the MFC in response to errors on these tasks (26), we focused on mediofrontal theta following errors (vs. correct) responses on an inhibitory control, Go/No-go task. Emerging work in developmental populations also suggests a possible dissociation between the mediofrontal theta response that immediately follows errors (early theta) compared to a relatively later response (late theta) (38). Thus, when comparing ASD and TD, we extracted error/correct mediofrontal theta from both an earlier and a later window and tested for possible dissociations. Finally, while the role of mediofrontal theta in laboratory-based cognitive control tasks is relatively well established (12), only limited work has sought to examine how these oscillations relate to more functional outcomes. Therefore, we also explored possible relations between early/late mediofrontal theta and academic and social outcomes. We hypothesized that mediofrontal theta would be reduced in children with ASD (compared to TD), and that such reductions would also be predictive of concurrent impairments in academic and social outcomes.

## Methods and Materials

### Participants

Participants included 43 children with ASD (M age=63.16 months, SD=4.30 months; 11 females) and 24 TD controls (M age=63.58 months, SD=4.86 months; 10 females) assessed at kindergarten-entry. The inclusion criteria were no cognitive delays (IQ≥85) and the regular use of complex sentences. Children with ASD were included if they had a previous diagnosis of ASD, which was confirmed with the gold-standard diagnostic measure, the Autism Diagnostic Observation Schedule-2, Module 3 (ADOS-2; 39). The ADOS-2 was administered by examiners who achieved research reliability, under the supervision of a licensed clinical psychologist. All children with ASD received scores in the ASD classification (comparison scores on the diagnostic algorithm ranging 4-10; 40). TD children were invited to participate in the study if they did not have any previous psychiatric or medical diagnoses. No TD children showed clinically elevated scores on the Child Behavior Checklist (CBCL; 41) externalizing or internalizing behaviors subscales. Children with ASD did show significantly elevated scores on CBCL externalizing and internalizing domains (*p*<0.001), thus these were controlled for in all subsequent analyses. No children were taking any psychotropic medications.

As shown in Table 1, independent samples *t*-tests revealed no significant differences between ASD and TD in nonverbal IQ (NVIQ) or age. The ASD sample included 53% White, 7% Black, 4% Asian, and 22% biracial children and 14% of other or unknown race. Similarly, the TD children were 43% White, 8% Black, 4% Asian, 34% biracial, and 11% of other or unknown race. A majority of caregivers (91% in TD and 85% in ASD) had at least a bachelor’s degree or higher. Most children were reported being right-handed (84% for ASD, 80% TD). All caregivers signed an IRB approved informed consent form.

**Table 1.** Sample Characteristics and Diagnostic Differences

|  | ASD | TD | ASD vs TD |
| --- | --- | --- | --- |
|  | *M* (SD) | *M* (SD) | Significance |
| *N* (females) | 43 (11) | 24 (10) |  |
| Age in months | 63.09 (4.27) | 63.48 (4.79) | n.s. |
| NVIQ | 106.14 (10.97) | 110.96 (10.17) | n.s. |
| CBCL Standard Scores |  |  |  |
| Internalizing Behaviors | 55.51 (11.70) | 43.33 (10.09) | <0.001 |
| Externalizing Behaviors | 53.51 (13.01) | 42.33 (9.56) | <0.001 |
| WJ Standard Scores |  |  |  |
| Applied Problems | 99.93 (14.63) | 108.00 (13.83) | 0.031 |
| Math Fluency | 86.60 (13.38) | 86.57 (8.80) | n.s. |
| Letter Word Identification | 108.12 (14.57) | 100.63 (11.42) | 0.034 |
| Passage Comprehension | 110.88 (13.86) | 107.24 (9.94) | n.s. |
| PIPPS T-Scores |  |  |  |
| Play Disruption* | 52.02 (9.97) | 48.23 (8.86) | n.s. |
| Play Disconnection* | 56.17 (16.43) | 30.57 (22.34) | <0.001 |
| Play Interaction<sup>#</sup> | 39.71 (12.96) | 54.35 (9.28) | <0.001 |

Note. Independent samples *t*-tests were employed to examine diagnostic differences.
\*Higher scores denote higher levels of play disruption and disconnection.
<sup>#</sup>Higher scores denote more play interaction.

### Behavioral Measures

#### Cognitive skills

The Differential Ability Scales (DAS; 42) was used to measure cognitive functioning for both groups. NVIQ was used as an estimate of cognitive ability in analyses, given that it is more stable than Verbal IQ in children with ASD (43). One child in the ASD and another in the TD group had NVIQs ± 3 SD from the mean were considered outliers and excluded from analyses.

#### Social Skills

The Penn Interactive Play Scale (PIPPS; 44) was used to measure social skills. This parent-report captures children’s play behaviors with their peers at home and in the community. It is comprised of three subdomains; Play Disruption, which assess aggressive play behaviors; Play Disconnection, which targets withdrawn play behaviors; and Play Interaction, which reflects play strengths. *T*-scores were used in analyses (*M*=50, SD=10). One TD child had a Play Disruption *t*-score ± 3 SD from the mean and was excluded from analyses. Additionally, 1 TD and 1 ASD children were missing scores for all domains.

### Academic achievement

Academic achievement was measured using Woodcock-Johnson III NU tests of achievement (WJ; 45). Math skills were captured by Applied Problems (math problem solving skills) and Math Facts Fluency (basic arithmetic skills) domains. Reading ability was captured by Passage Comprehension (understanding of written text) and Letter-Word Identification domains. All subtests yield standard scores (*M*=100, SD=15), which were used in statistical models. Outliers included 1 child with ASD for Math Facts Fluency and 1 child with ASD for Passage Comprehension who received scores ±3 SD from the mean in each subdomain.

### Electrophysiological Tasks and Measures

#### EEG/Event-Related Potential (ERP) Task

EEG recordings were collected while children played a child-friendly Go/No-go task (Zoo Game; 46,47) in a testing room with minimal distractions. The Zoo Game successfully elicits EEG activity associated with error monitoring in TD children as young as 3 years (46). Children are instructed that they are playing a game to help a zookeeper catch the loose animals that escaped their cages in the zoo. To catch an animal, children are told to click an identified button when a picture of a loose animal appears (Go trials) and they are not supposed to press the button when they see any of the three orangutans who are helping them (No-go trials). Children start the game with a practice block. The actual task includes 8 blocks of 40 trials (320 trials total; 240 Go and 80 No-go). The stimuli are presented for 750 ms and then a blank screen for 500 ms. All images are preceded by a fixation cross randomly jittered between 200-300 ms. Children can make their responses while the stimulus is on the screen or during the 500 ms blank screen that follows (1250 ms response deadline). Feedback on performance is provided to children after each block with prompts generated based on the calculation of error rates to ensure an acceptable number of trials for stable EEG waveforms. No trial-level feedback was provided. Zoo Game behavioral accuracy was captured by the percentage of error/correct trials for Go and No-go trials. Response time (RT) was also assessed for overall, correct-Go, and error-No-go trials.

### Electrophysiological recording, data reduction, and data processing

#### EEG Recording

Stimuli from the Zoo Game was presented on a PC laptop using E-Prime 2.0. Net Station 5.4, running on a Macintosh laptop was used to record EEG from a 64-channel Geodesic sensor net (EGI). Impedances for all electrodes were kept below 50 kΩ, following recommendations for this system. The EEG signal was digitized and sampled at 500 Hz via a preamplifier system (EGI Geodesic NA 400 System).

#### EEG preprocessing

EEG data were processed using Matlab (The MathWorks, Natick, MA), EEGLAB (48), FASTER (49), ADJUST (50) and custom Matlab scripts partly based on work by Bernat and colleagues (51). Preprocessing methods reflect a precursor to the Maryland Analysis of Developmental EEG (MADE) pipeline (52). Briefly, EEG data were digitally filtered and bad channels removed. A copy of the dataset was further cleaned via automated methods prior to running ICA. ICA weights were copied back to the original dataset and used to remove ocular and other artifacts (53,54). Data were epoched to response markers from −1000 to 2000 ms and baseline corrected using the −400 to −200 ms pre-response period. A final rejection of residual ocular artifacts using a +/-125 µV threshold was conducted, missing channels were interpolated, and an average reference was computed. See supplement for complete details.

#### Time-frequency decomposition

Given our focus on theta (∼4-7 Hz) oscillations, the EEG data were down-sampled to 32 Hz to improve computational efficiency with no loss of the signals of interest (i.e., Nyquist = 16 Hz). Cohen’s class reduced interference distributions (RIDs) were used to decompose time frequency (TF) representations of response-locked power for each trial before averaging across all trials (51). TF surfaces were baseline-corrected relative to the −400 to −200 (pre-response) period using a subtractive baseline correction to isolate response-related changes in power. Data were not converted to a dB scale and units reflect response-related changes in (baseline-corrected) raw power. As part of a standard preprocessing pipeline of TF data that allows meeting additional assumptions necessary for synchrony-based analyses, a subsampling approach was implemented (10). However, given that the subsampling procedure was conducted only as a preprocessing step for analyses of synchrony (not reported here) and do not serve to improve the reported power-based analyses, details of the subsampling procedure are described elsewhere (10).

#### Extraction of response-related frontal theta power

Given prior work in adults (28), adolescents (11), and children (37) linking error-related changes in theta power over frontal scalp sites to error monitoring and cognitive control, we extracted response-related theta power (4-7 Hz) from a cluster of three mediofrontal electrodes (E4/FCz, E7, E54). In line with prior work, we extracted frontal theta power during a predefined ROI based on the first ∼200 ms following the response (exact window timing based on sampling resolution; first 6 samples = 0-192 ms).

Consistent with emerging work suggesting a dissociation between “early” and “late” theta power following responses, we additionally extracted response-related theta power from a second ∼200 ms window immediately following our first ROI (exact window timing based on sampling resolution; next 6 samples = 192-384 ms). Thus, for each participant, response-related theta power within the 4-7 Hz band was extracted from a frontocentral cluster of electrodes during an early (first ∼200 ms) and late (next ∼200 ms) window, separately for error and correct trials.

Determining how many clean EEG trials each participant should have, in order to be included in error-related analyses, involves balancing reliability of the EEG signal on the one hand, and risk of creating a biased sample on the other. Creating a biased sample through participant exclusion is particularly problematic within the context of comparisons between clinical and non-clinical groups. Moreover, simulation studies have yet to identify the optimal number of trials necessary for calculating reliable error-related theta signals. However, at least one simulation study of the associated “error-related negativity” (ERN), a time domain EEG signal that theta is known to contribute to substantially (55), suggests that 4-6 trials may be sufficient for computation of a reliable ERN (56). Therefore, we conducted analyses of the time-frequency data utilizing an inclusion criterion of 4 trials (and a minimum Go accuracy of 50%) to maximize participant inclusion and guard against sample biases.

### Statistical Analyses

Independent *t*-tests were conducted to examine differences in social and academic skills between ASD and TD. Generalized Linear Mixed Models (GLMMs) were conducted to examine diagnostic differences (ASD vs TD) in accuracy and reaction time (RT) on the Zoo Game. RT data are known to be positively skewed (57), therefore analyses were conducted with log-transformed RT. At the neural level, a GLMM with a three-way interaction term (accuracy [correct vs. error] by timing [early vs. late theta] by diagnosis [ASD vs. TD]) was used to examine whether children exhibited error-related theta (significant increases in theta power for error vs. correct trials) and whether these effects differed as a function of diagnosis or timing (early vs. late theta ROIs). Post-hoc analyses explored the nature of any significant interactions. Lastly, regression analyses explored whether error-related differences in early or late theta power predicted academic and social skills across both groups. Age, gender, internalizing and externalizing domain *t*-scores on CBCL, and NVIQ were controlled for in all GLMMs and regressions. All analyses were conducted using SPSS Version 24.

## Results

### Diagnostic Differences in Social and Academic Skills

Based on parent-report, children with ASD showed significantly more impairments in Play Interaction (*t=*-4.78, *p*<0.001) and Disconnection (*t*=5.27, *p*<0.001) compared to TD. In areas of academic achievement, the ASD group scored significantly lower than the TD group in math (Applied Problems; *t*=2.21, *p*=0.031). Children with ASD scored significantly higher than the TD sample for Letter Word Identification (*t*=-2.17, *p*=0.034). See Table 1.

### Diagnostic Differences in Zoo Game Performance

Diagnosis emerged as a significant main effect, such that children with ASD performed worse than the TD sample for Zoo Game accuracy (*F*=17.76, *p*<0.001), while controlling for age, gender, internalizing and externalizing behaviors, and NVIQ. There was also a main effect of trial type, with accuracy being lower on No-go trials than Go trials for all children (*F*=95.76, *p*<0.001). However, there was no significant interaction effect for diagnosis and trial type accuracy. In regard to RT, there was a main effect of trial type, with all children showing slower RT for correct responses on Go trials compared to error responses during No-go trials (*F*=83.14, *p*<0.001). No significant diagnostic differences were observed in RT. Results using raw RT were similar. See Table 2 for details on accuracy and RT by diagnostic group.

**Table 2.**
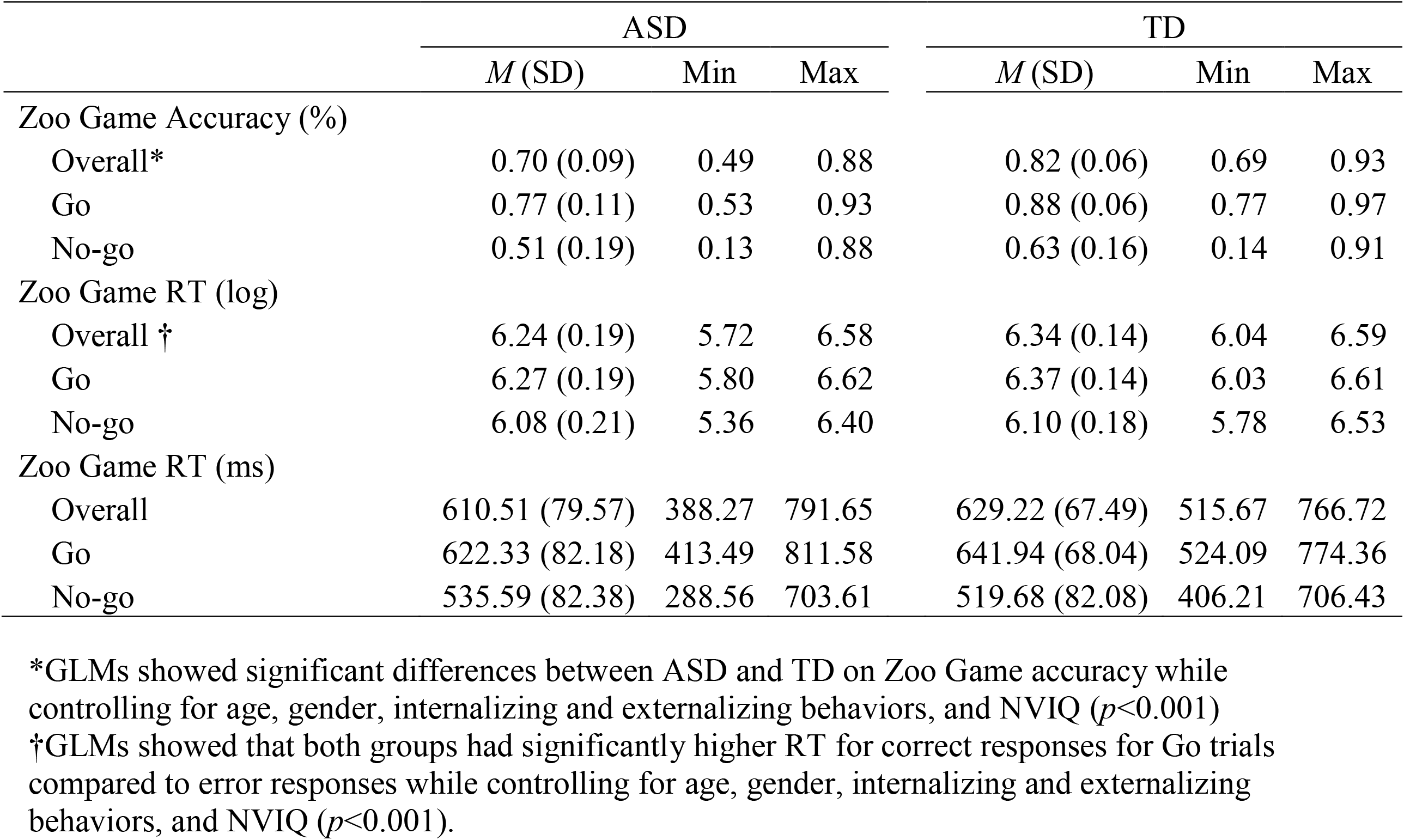
Behavioral Performance on the Zoo Game by Diagnostic Group

The Presence of Error Related Theta Power and Diagnostic Comparisons on Early vs. Late Theta A GLMM revealed a significant main effect of accuracy (*F=*9.81, *p*=0.002), while controlling for age, gender, internalizing and externalizing behaviors, and NVIQ, confirming higher mediofrontal theta power for error vs. correct trials overall (Figure 1). A significant main effect of diagnosis (*F*=14.63, *p*<0.001) emerged, with the ASD group having overall lower mediofrontal theta power. Crucially, there was also a significant three-way interaction between accuracy, timing of theta, and diagnostic group (*F*=3.56, *p*=0.030). To explore the nature of this interaction, a pair of post-hoc, two-way (accuracy [correct vs. error-trials] by diagnosis [ASD vs. TD]) GLMMs were conducted to explore specific diagnostic differences in theta between error and correct trials for each theta timing ROI. A significant two-way interaction of accuracy x diagnosis was revealed for the late theta ROI (*F*=4.82, *p*=0.030), such that TD children exhibited a larger error (vs. correct) increase in late theta power relative to children with ASD (Figure 2). In contrast, no interaction between accuracy and diagnosis emerged for the early theta ROI (*F*=2.00, *p*=0.160).

**Figure 1.**
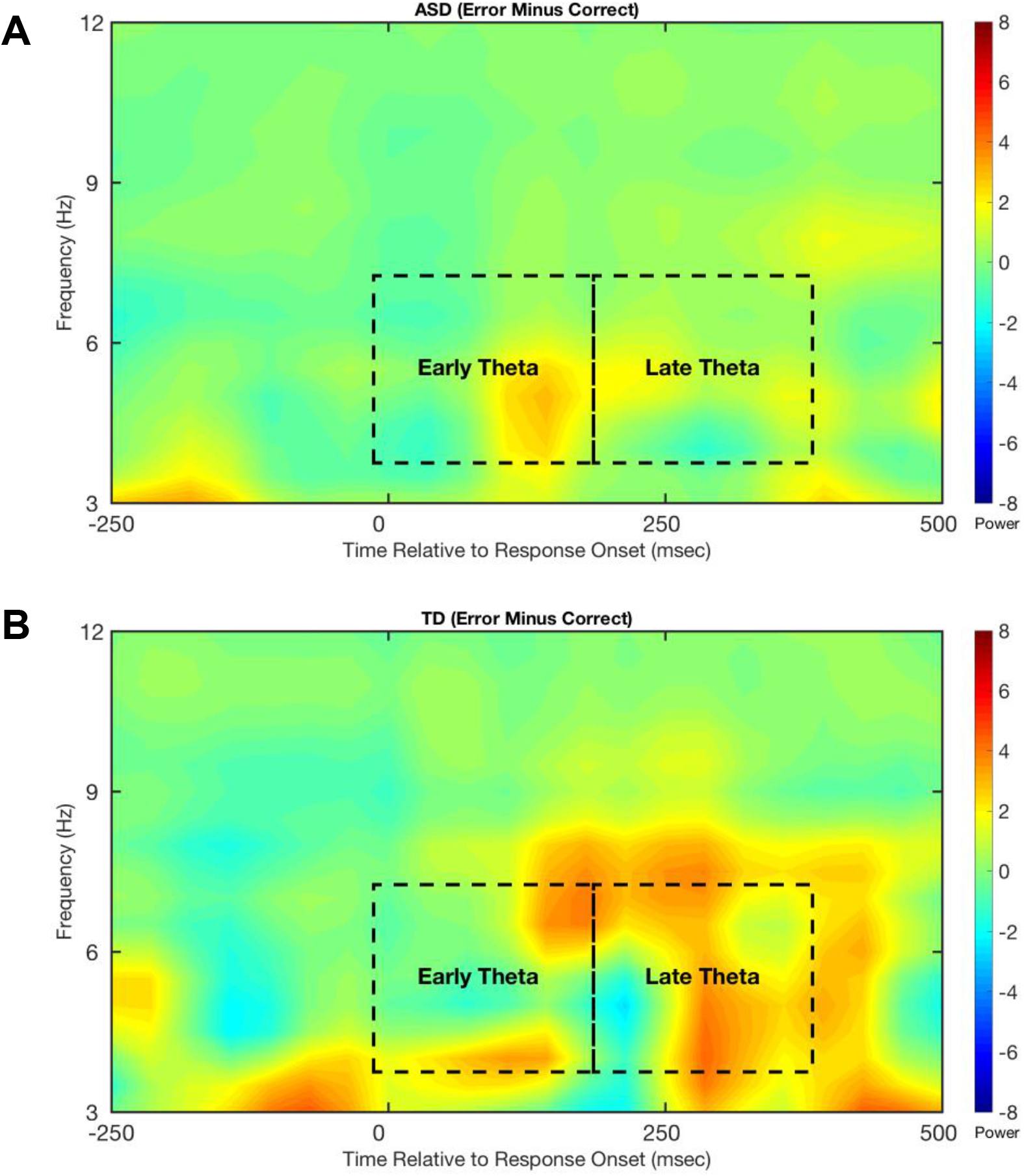
Time-frequency plots of response-locked error-related (error minus correct) power at a mediofrontal electrode location (E4/FCz). Zero msec corresponds to the time of response commission. Units of power reflect baseline-corrected raw power. The regions theta-band (4-7 Hz) regions of interest for “early” and “late” theta are indicated with dashed boxes. Error-related time-frequency surfaces are plotted separately as a function of diagnostic group: A) children with autism spectrum disorder (ASD); B) typically developing children (TD).

**Figure 2.**
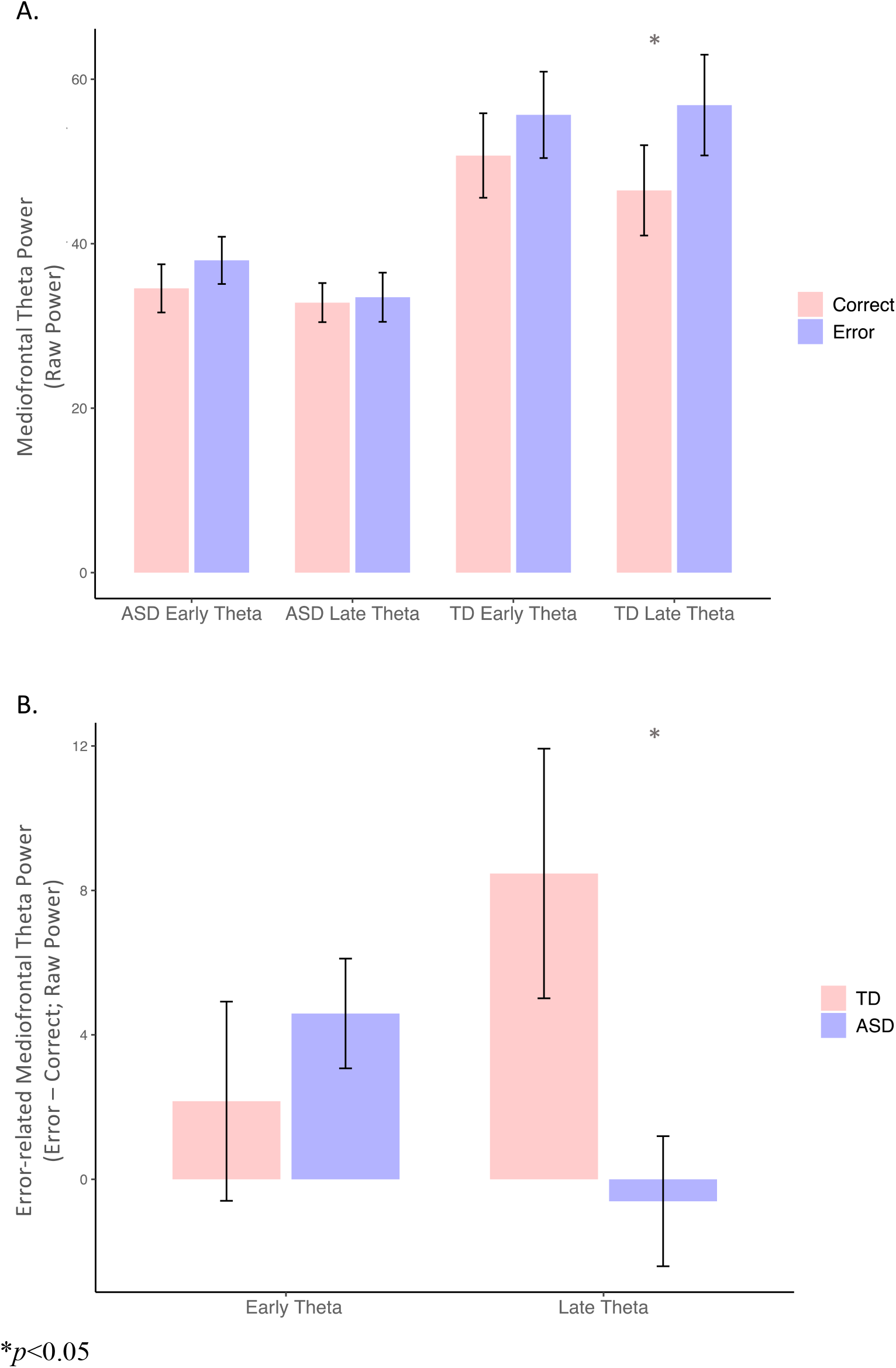
A) Diagnostic differences in early and late mediofrontal theta power by response type; B) Comparison of early and late window accuracy differences in mediofrontal theta power between diagnostic groups.

### Theta Power Predicting Social Skills and Academic Achievement

Regression analyses showed that late theta power significantly predicted math skills on the WJ Applied Problems subdomain (*p*=0.01) and social skills on the PIPPS Play Interaction subdomain (*p*=0.045), while controlling for age, gender, internalizing and externalizing behaviors, and NVIQ. See Table 3 for detailed results of the significant regression results. Early theta did *not* significantly predict any of the social or academic domain measures (all *p*>0.05).

**Table 3.**
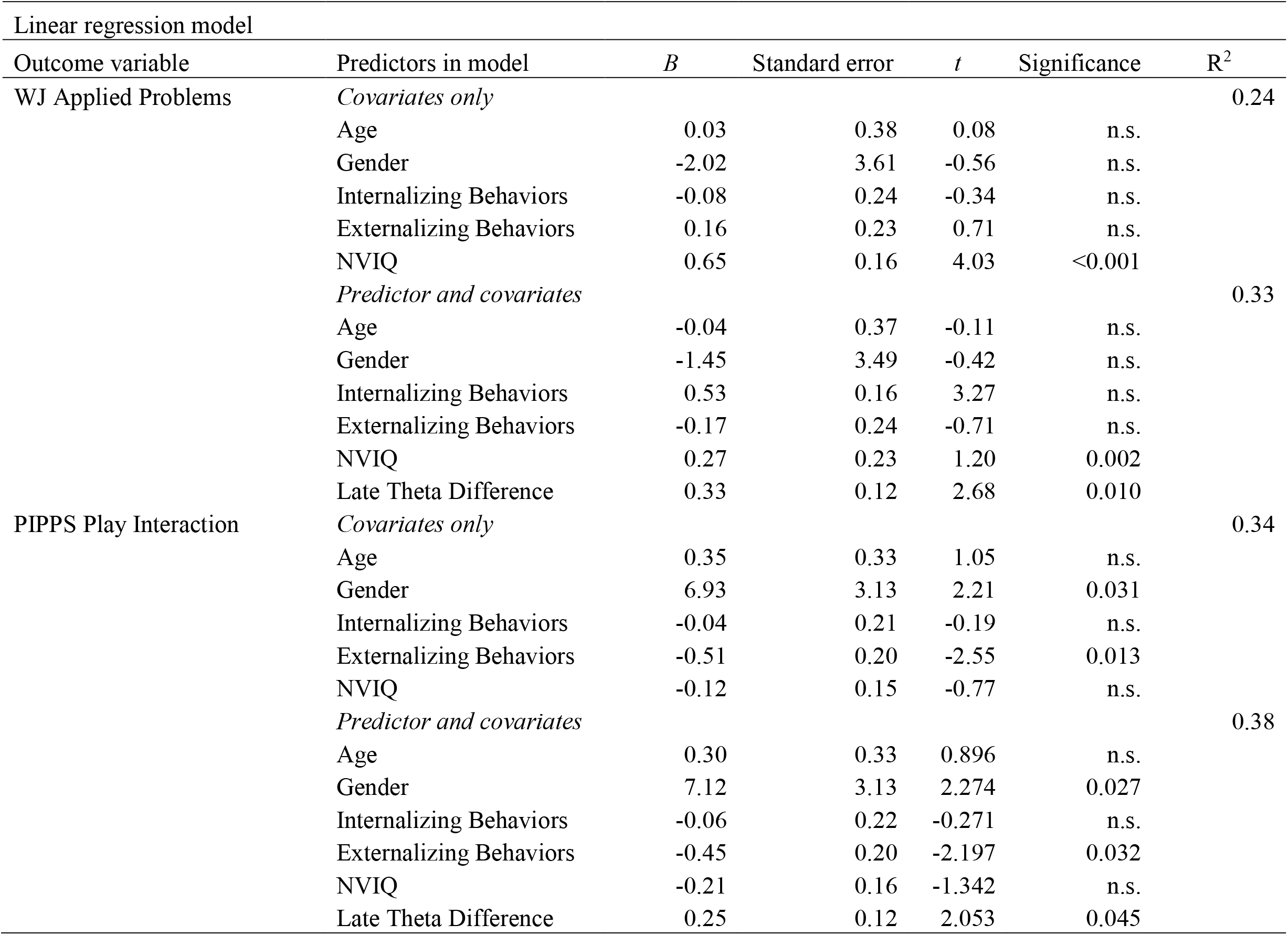
Regression of late theta power as a predictor for academic and social skills for the combined ASD and TD sample

## Discussion

The current study is one of the first to demonstrate that task-related mediofrontal theta oscillations underlying cognitive control are disrupted in kindergarteners with ASD compared to TD. While individuals with ASD did not exhibit significant differences in theta immediately following error-responses (early theta), they displayed a reduced magnitude in later emerging mediofrontal theta (late theta). Moreover, these differences in late, but not early, theta predicted academic and social outcomes. These data are consistent with prior work reporting impairments in cognitive control in ASD at the behavioral level (7,20). Similarly, the current data are in line with results from neuroimaging studies that identify functional (23,24,26) and structural (26,27,58) abnormalities associated with the MFC and cognitive control networks in ASD. However, EEG provides the novel opportunity to directly assess oscillatory activity not captured by behavioral or neuroimaging (f*/*MRI) approaches. Given that mediofrontal theta oscillations are causally linked to cognitive control (13,14) and EEG affords the opportunity to more precisely identify transient brain activty, our findings may provide direct insight into understanding *how* and *when* cognitive control is disrupted in young children with ASD. Of equal importance, the current report grounds the assessment of mediofrontal theta oscillations in terms of how they relate to more traditional functional outcomes (social and academic domains).

Utilizing a child-friendly Go/No-go task (Zoo Game; 46), we extracted error/correct mediofrontal theta power measures in kindergarteners with ASD and typical peers to examine behavioral and neural differences in cognitive control. Behaviorally, children with ASD showed reduced overall accuracy on the Zoo Game, even after accounting for age, gender, internalizing and externalizing behaviors, and NVIQ. These results replicate studies indicating behavioral deficits in tasks targeting general cognitive control in this clinical population (7,20,59). However, task-related behavioral measures did not reveal differences in more specific EF domains, such as changes in overall attention (i.e., Go accuracy) or inhibitory control (i.e., No-go accuracy). Thus, relying solely on behavioral measures may limit a more detailed examination and nuanced understanding of how and why individuals with ASD exhibit deficits in cognitive control. In contrast, employing analyses of mediofrontal theta in the current study allowed a more comprehensive assessment of cognitive control dynamics within ASD.

To further investigate the neural underpinnings of impairments in cognitive control in ASD, we compared response-related mediofrontal theta oscillations between ASD and TD children based on their *magnitude* and *timing*. Drawing on work suggesting possible dissociations between mediofrontal theta oscillations that arise relatively earlier or later in the post-error period (38), we separately analyzed an “early theta” (∼0-200 ms) and “late theta” (∼200-400 ms) time window. Notably, children with ASD (compared to TD) exhibited a selective reduction in mediofrontal theta power during the late, but *not* early, time window. Similarly, a recent study examining event-related theta power in older children with ASD also found selective reductions theta within a later post-event time window when performing a cognitive flexibility task that relies on higher-order cognitive processes such as inhibition and attention shifting (60). Cognitive control is generally thought to involve a cascade of processing whereby the need for control is first detected and later followed by the recruitment and allocation of top-down control to bias behavior favorably (1). Thus, error-related theta power within the early time window may be associated with the detection of errors and need for control, whereas the late theta time window may more closely map onto higher-level neural processes associated with the recruitment of top-down control. The current data suggest that this second stage of cognitive control may be particularly impaired in young children with ASD compared to TD.

Consistent with the notion that late theta may more closely relate to higher-level aspects of cognitive control, we found that increased error-related theta power during the late time window, but *not* early theta, was a significant predictor of academic (math) and social skills in ASD and TD children. Various behavioral studies have shown that impairments in cognitive control may negatively impact academic and social functioning (7,15-19). Moreover, our previous work found that variability in mediofrontal theta predicted math abilities in children with ASD (37). The current study further extends these results by revealing a more specfic pattern of neural dyanamics that may play a key role in social and academic development in young children.

### Limitations and Future Directions

Given the heterogeneity of behavioral presentation in children with ASD, and developmental effects on cogntive control, the current study included a focused sample of verbal, kindergarten-aged children with ASD without cognitive delays (IQ≥85). Therefore, results need to be replicated with a larger sample across a wider range of ages and abilities. TD children had no previous psychiatric diagnoses or clinically elevated CBCL externalizing or internalizing scores, however, children with ASD did show elevated scores. While we statistically controlled for externalizing and internalizing behaviors, future work should explore ASD and TD children matched for levels of externalizing and internalizing problems to examine the inter-relations among these co-occurring symptoms and cognitive control.

## Conclusions

Prior work finds that individuals with ASD have deficits in cognitive control at the behavioral level and neurally exhibit structural and functional abnormalities within the MFC. However, previous studies have not directly investigated whether task-related mediofrontal theta oscillations, thought to arise from the MFC, are disrupted in young children with ASD. We identified selective reductions in error-related theta oscillations emerging in a reltively late post-error time window. Moreover, such reductions in late theta were found to predict academic and social outcomes. These results indicate that accounting for the neural dynamics of cognitive control in individuals with ASD may provide specific information about deficits not observable behaviorally. Moreover, the results, if replicated, suggest a novel neurocognitive target for interventions designed to support social and academic outcomes in cognitively-able children with ASD.

## Acknowledgements

This project was funded by the Kellen Junior Faculty Award (PI: Kim). We thank Catherine Lord, Nathan Fox, Frederick Morrison, Jennifer Grammer, William Fifer, Natalie Brito, Lauren Shuffrey, Susan Faja, Nurit Benrey, Claire Klein, and Denisse Janvier for help with data collection. We thank the children and the families who participated in the study.

