## Supplement for "Atypical mediofrontal theta oscillations underlying cognitive control in kindergarteners with autism spectrum disorder"

**EEG preprocessing**

EEG data were processed using Matlab (The Mathwoks, Natick, MA), the EEGLAB toolbox (40), the FASTER toolbox (46), the ADJUST toolbox (48) and custom Matlab scripts partly based on work by Bernat and colleagues (42). Preprocessing methods reflect a precursor to the Maryland Analysis of Developmental EEG (MADE) pipeline, which is described in detail elsewhere (46). See supplement for complete details of EEG preprocessing. Breifly, EEG data was digitally filtered offline (0.3 Hz high-pass, 50 Hz low-pass) and globally bad channels were detected and removed using the FASTER toolbox (47). In order to classify a channel as artifactual, FASTER calculates three parameters: variance, mean correlation, and Hurst exponent. Channels with a Z-score of ±3 for any parameter were deemed to be globally bad and were removed from further analyses. In preparation for independent component analysis (ICA), a copy of the original dataset was created and then filtered using a 1 Hz high-pass filter (41,43). This copied data set was segmented into arbitrary 1 second epochs to identify and remove epochs with excessive artifact; artifacts were detected and removed if amplitude was +/- 1000 uV or if power within the 20-40hz band (after Fourier analysis) was greater than 30dB. ICA was then performed on the copied, high-pass filtered dataset after removing epochs with artifacts. Following ICA decomposition, ICA weights were copied back to the original dataset (41,43); components associated with ocular or other artifacts were identified and removed. The data were then epoched to the response markers from -1000 to 2000 ms and baseline corrected in the time domain using the -400 to -200 ms period preceding the response. Epochs with residual ocular artifacts were identified and removed using a +/- 125 µV threshold based on the ocular channels. For all remaining channels, for epochs in which a given channel exhibited voltage +/-125 µV, data were removed and interpolated using a spherical spine interpolation, unless more than 10% of channels were bad within a given epoch, in which case the entire epoch was removed. Finally, any channels that were marked as bad throughout the entire recording were interpolated (spherical spline interpolation) and an average reference was computed.
